# FARM: Forecasting Antibiotic Resistance in *Mycobacterium tuberculosis* using biophysics and machine learning

**DOI:** 10.64898/2026.07.23.740359

**Authors:** Mahbuba Tasmin, Shrishti Barethiya, Yu Wang, Lulu Kang, Jianhan Chen, Anna G. Green

**Affiliations:** Manning College of Information and Computer Sciences, University of Massachusetts, Amherst, MA 01003; Department of Chemistry, University of Massachusetts, Amherst, MA 01003; Department of Mathematics and Statistics, University of Massachusetts, Amherst, MA 01003

## Abstract

Antibiotic-resistant tuberculosis remains a major public health challenge, and rapid diagnosis of resistant infections based on genomic markers holds promise for improving time to effective treatment. However, the vast majority of clinically observed variants in resistance-associated genes remain of uncertain significance, limiting the utility of predictors. Here we develop a multimodal forecasting framework, FARM (Forecasting Antibiotic Resistance in *Mycobacterium tuberculosis*) to determine whether a newly observed mutation in a resistance gene may indeed cause resistance. Our framework combines structural context, biophysical energy features, protein language model features, and mutational AAIndex physicochemical descriptors. Using 345 labeled mutations from the World Health Organization 2021 catalogue, we train interpretable models that distinguish resistance-associated from non-resistance-associated variants with holdout AUCs of 0.843–0.943. In a novel temporal evaluation of 62 mutations reclassified after the training data was released, the selected Combined model achieved 80.7% recall of resistant reclassifications (resistant-class F1=86.8; AUC=0.735). Applied to 4,525 current uncertain-significance mutations, the framework prioritizes 696 candidate resistance mutations, including genes associated with the new antibiotics bedaquiline, delamanid, and pretomanid. These forecasts are intended to support future catalogue updates and experimental follow-up.

## 1 Introduction

Tuberculosis (TB) remains one of the leading causes of infectious disease mortality worldwide, and the rise of drug-resistant strains has made treatment increasingly challenging [1, 2] Determining an effective antibiotic regimen depends on drug susceptibility testing (DST), yet phenotypic DST for *Mycobacterium tuberculosis* requires 3–6 weeks [3, 4] During this delay, patients may receive ineffective or suboptimal treatment, worsening disease outcomes and enabling further transmission and evolution of resistance. Rapid and accurate resistance profiling at the time of diagnosis is therefore essential for improving clinical decision-making and global TB control.

Genomic sequencing provides a promising alternative to phenotypic DST, enabling genotype-based prediction of drug resistance directly from bacterial DNA. Rule-based diagnostic approaches built around curated resistance mutation catalogues, such as those maintained by the World Health Organization (WHO), perform well for well-characterized variants, particularly for first-line drugs [5–9]. However, performance drops for second-line and newer drugs, where mechanisms are more heterogeneous and catalogue coverage remains incomplete. As a result, many clinically observed mutations remain classified as having “uncertain significance.”

Computational approaches for identifying resistance-conferring mutations in *M. tuberculosis* broadly fall into two paradigms: genotype-to-phenotype prediction and variant-to-effect prediction. In genotype-to-phenotype prediction, models are trained on individual isolate genome sequence features to predict isolate phenotypes, and *post hoc* interpretation is used to identify contributing variants. These approaches perform well for several first-line drugs but generalize less well to second-line and newer regimens, where mechanisms are more heterogeneous and labeled data are scarcer [10–18]. Because these models rely largely on sequence-derived features from sequenced and phenotyped isolates, they can struggle to interpret rare or previously unseen mutations, which remain abundant in the current WHO catalogue.

In contrast, variant-to-effect prediction uses a specific genetic variant as input and predicts its functional impact from features of that variant. This formulation is particularly well suited to *M. tuberculosis*, which acquires resistance primarily through chromosomal point mutations [19–21]. Prior studies in this paradigm have largely focused on individual gene–drug systems using structural and physicochemical mutation descriptors [22–27], though recent works have extended to multiple gene–drug combinations in a single model [28, 29]. Related work in viral systems has shown that combining structural and evolutionary information can improve forecasting of mutation effects [30–33].

In this work, we introduce the FARM framework for forecasting whether non-synonymous variants designated Uncertain Significance in the WHO catalogue do indeed lead to antibiotic resistance (Fig. 1). We reason that such variants may be distinguished from neutral variants by their shared evolutionary, structural, and biochemical features, as has been successful in past work [22–27, 29, 28]. We build a classification system that uses 25 multimodal features to predict whether a given variant is resistance-associated. When trained on known-effect variants from the 2021 WHO catalogue, the best-performing models achieve holdout AUCs of 0.843–0.943 across gene-regime settings. We then use a temporal evaluation setup to assess performance on variants newly classified in 2023, after the release of the training data, where the deployed model recovers 80.7% of resistant reclassifications with resistant-class F1 of 86.8%. We further show that the framework learns biologically interpretable features associated with resistance prediction. Finally, we provide a prospective classification of 4,525 latest uncertain-significance variants from WHO 2023 catalogue, including variants in genes associated with resistance to newly introduced drugs. These forecasts provide a practical prioritization resource for experimental validation and future catalogue updates.

**Figure 1:**
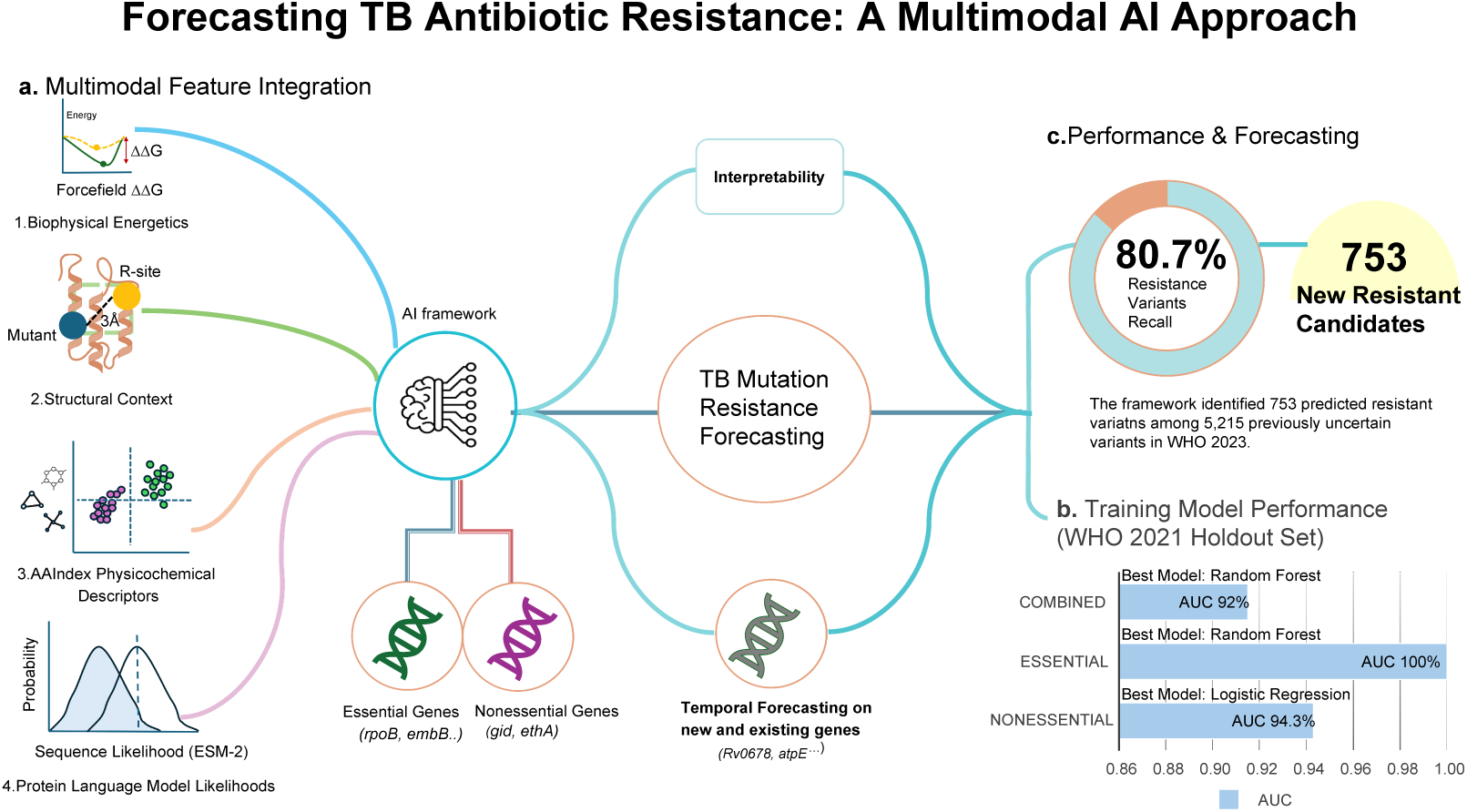
FARM: A multimodal forecasting framework for *M. tuberculosis* resistance. **(a) Feature integration:** each amino-acid substitution is represented by 25 features spanning structural context, Biophysical energy terms, ESM-2 sequence-likelihood features, and AAIndex physicochemical descriptors.**(b) Model selection:** models trained on WHO 2021 labeled variants capture distinct gene-class regimes, with random forests performing best for Essential and Combined sets and logistic regression performing best for the Nonessential set.**(c) Temporal forecasting:** the final framework is evaluated on WHO 2023 reclassifications and then deployed to forecast uncertain-significance variants, including genes with no direct 2021 labeled support.

## 2 Results

### 2.1 Mutation-level cohort curation for antibiotic resistance forecasting

We seek to build a method that can, given a genetic variant observed in a resistance gene in *M. tuberculosis*, predict whether that variant is associated with antibiotic resistance. For training data, we use the catalogue of known resistance-conferring mutations published by the World Health Organization in 2021[34, 6]. Variants in this catalogue are designated as known to be associated with resistance, known not to be associated with resistance (consistent with susceptibility), or variants designated Uncertain Significance. Thus, it serves as an ideal dataset for training and evaluating our model.

Starting from the WHO 2021 catalogue, we restricted the analysis to non-synonymous substitutions and retained unique feature-complete gene-mutation entries after quality control (Fig. 2). This produced 345 known-effect mutations used for supervised training and 3,412 uncertain-significance mutations excluded from training. Of these uncertain-significance mutations, 62 were reassigned to known-effect categories in the 2023 WHO catalogue, forming the temporally separated evaluation set. We excluded synonymous, insertion/deletion, and non-coding variants from this analysis. While these mutation classes also contribute to resistance, including through gene regulatory effects, they would require a different feature representation from the protein-centered multimodal framework used here.

**Figure 2:**
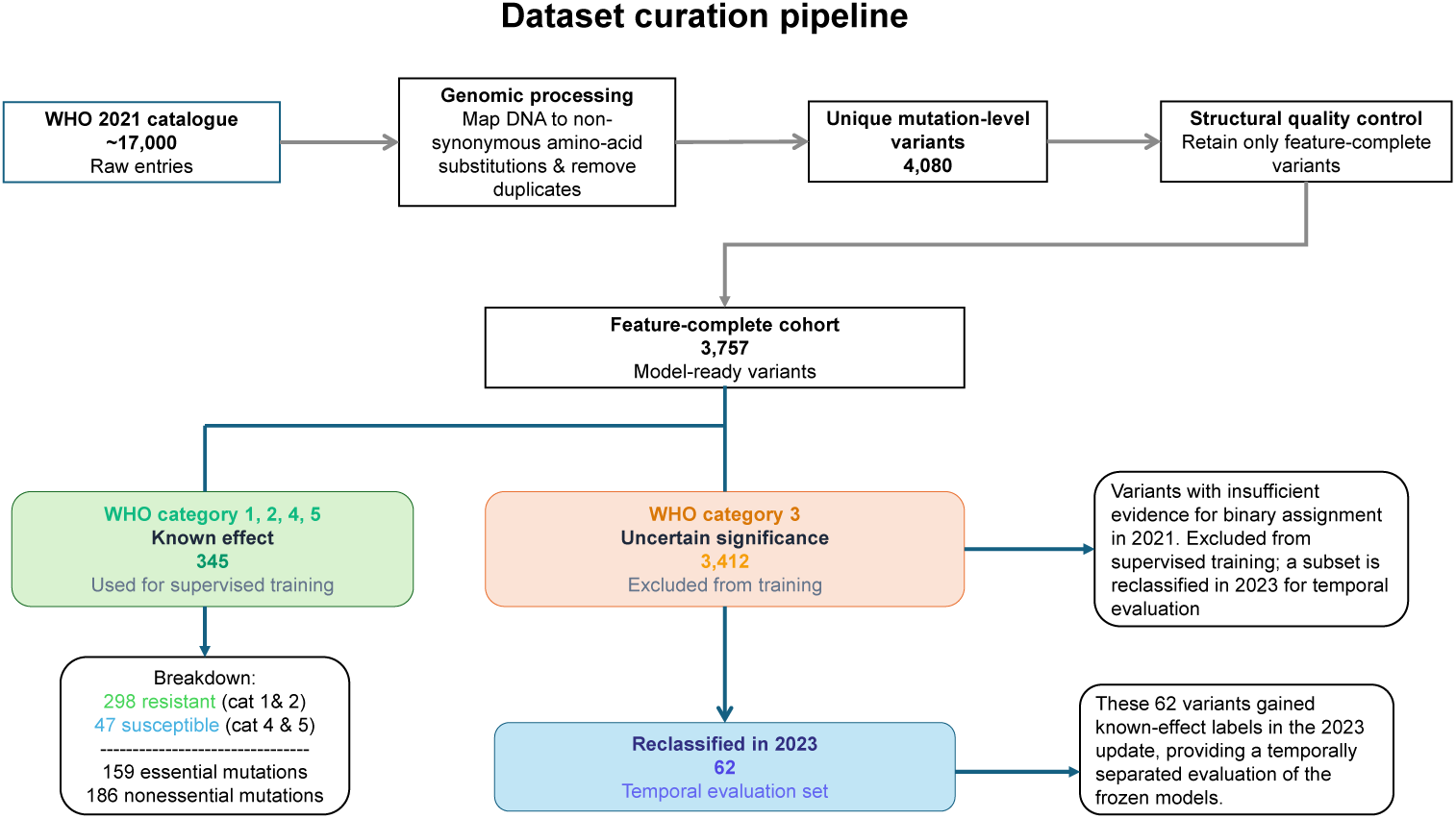
Dataset curation and mutation-level cohort definition for resistance forecasting. Workflow for deriving the 2021 mutation-level training cohort and temporally separated evaluation set from the WHO catalogue.

For each variant, we computed a 25-feature multimodal representation comprising structural context features derived from protein 3D structures, biophysical energy features derived from Rosetta-based structural modeling, physicochemical descriptors of amino acids from AAIndex, and protein language model features from ESM-2, as summarized in Table 1 and described in Section 4.1 and Section 4.2. These features served as the input representation for predicting whether a variant would be labeled as resistance-associated or not resistance-associated. We also stratified the analysis into variants assigned to the Essential and Nonessential gene strata, reasoning that these proteins are subject to different *in vivo* functional constraints and may therefore exhibit distinct mutational regimes associated with resistance (see Section 4.1). We term these models the Essential and Nonessential models, in contrast with the Combined model trained on all labeled training mutations.

**Table 1:** Feature groups in the 25-feature multimodal representation used for training, temporal evaluation, and forecasting.

| Feature group | # | Biological interpretation |
| --- | --- | --- |
| Structural context | 1 | Distance from the mutated residue to known resistance-associated positions within the same gene, used as a structural anchor signal. |
| Biophysical energy | 6 | Energetic and stability consequences of the mutation from Rosetta-based structural modeling. |
| Protein language model features | 10 | Sequence-context plausibility from the ESM-2 language model, represented as selected latent coordinates rather than direct mechanistic variables. |
| AAIndex physicochemical descriptors | 8 | Physicochemical summary of the mutant amino acid in AAIndex space. |

### 2.2 Model training on labeled WHO 2021 variants

We trained binary classifiers to predict, for a given input variant described by its multimodal features, whether that variant would be labeled as resistance-associated. We compared logistic regression and random forest architectures on a label-stratified 80/20 holdout split of the 2021 WHO catalogue, with model selection and threshold determination performed via cross-validation within the training data as described in Section 4.3, 4.4. Under this procedure, random forest was the best-performing architecture for the Combined setting and achieved a holdout AUC of 0.920 on the unseen test split, with resistant-class F1 of 95% (Fig. 3A; Supplementary Table S5).

**Figure 3:**
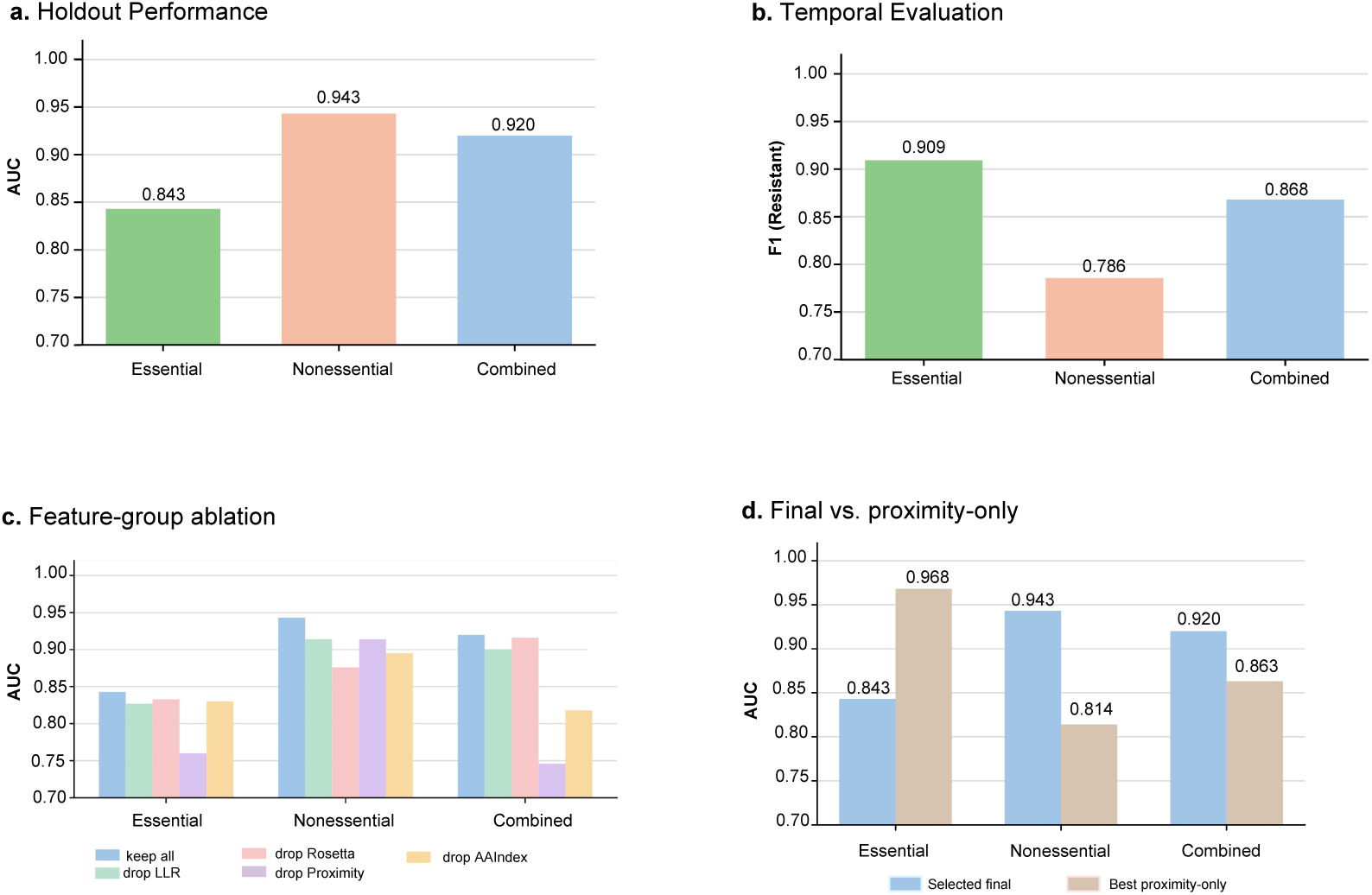
Predictive performance and feature-group dependence across gene strata. (a) Holdout AUC for the best-performing model in each dataset configuration. (b) Temporal resistant-class F1 on the subset of 2021 uncertain-significance variants reclassified by WHO in 2023. (c) Single-group ablations for the selected best model in each dataset highlight distinct dependence on structural context versus protein language model features. (d) Comparison between the selected final model and the best structural context-only baseline in each dataset.

We then tested whether splitting the data into variants found in the Essential and Nonessential genes, and retraining models separately, could improve performance by capturing differences in underlying biological signal. Random forest performed best for the Essential setting, achieving holdout AUC 0.843 and resistant-class F1 92.6% on a test set of 37 variants. Logistic regression performed best for the Nonessential setting, achieving holdout AUC 0.943 and resistant-class F1 90.6% on a test set of 38 variants. Thus, splitting by gene stratum improved holdout performance in the Nonessential setting but not in the Essential setting.

### 2.3 Ablation study: contribution of feature groups

To identify which of the multimodal features drive discrimination between resistance-associated and non-resistance-associated variants, we performed an ablation study to examine the importance of the four main feature groups: protein language model features, biophysical energy features, structural context, and AAIndex physicochemical descriptors. Figure 3 summarizes single-group removals for the best-performing model in each dataset, with full grids reported in Supplementary Table S7. Across datasets, structural context was the most consistently critical group, followed by biophysical energy features and protein language model features. Removing structural context caused the largest performance loss in Essential RF (AUC 1.00 *→* 0.78) and Combined RF (AUC 0.92 *→* 0.79).

In contrast, Nonessential Logistic was most sensitive to removal of protein language model features (AUC 0.94 *→* 0.91, sensitivity 0.83 *→* 0.69) and, secondarily, biophysical energy. These results suggest that the predictions from the Essential RF and Combined RF models rely most on structural anchoring to known resistance-conferring sites, with substitutions near binding, catalytic, or interaction interfaces contributing strongly to discrimination in these settings. In contrast, the Nonessential model predictions depend more on sequence-derived signal, possibly because resistance-associated mutations in the Nonessential genes more often reflect loss-of-function events that are distributed throughout the protein. Structural-context-only baselines are summarized in Fig. 3D and Supplementary Table S6.

In the Essential setting, the structural-context-only baseline outperforms the multimodal model, consistent with the dominant effect of structural context in this regime. In the Combined setting, the best structural-context-only baseline remained informative but did not exceed the full multimodal model, indicating that the additional feature groups contribute useful signal beyond structural context alone. We therefore retained the full Combined model as the main deployment model for forecasting the 2023 uncertain-significance variant set (Section 2.7) and interpret the single-feature baseline results as evidence that structural context is a particularly strong component of prediction in the Combined regime.

### 2.4 Temporal evaluation on 2021 uncertain-significance variants reclassified by WHO in 2023

A critical application of FARM is forecasting whether newly observed uncertain-significance mutations are likely to be associated with antibiotic resistance. To evaluate this temporal generalization setting, we applied models trained on known-effect mutations from the 2021 WHO catalogue to the 2021 uncertain-significance variants pool. Among these, 62 mutations were reassigned to known-effect categories in the 2023 WHO catalogue, providing a temporally separated evaluation set. Each model used decision thresholds fixed during training on the 2021 dataset (Section 4.3). Most reclassified mutations belonged to the Nonessential gene set (55 of 62 total).

Because this temporally separated evaluation set is imbalanced and the main biological question is recovery of resistant reclassifications, we summarize temporal evaluation performance primarily with resistant-class F1, while retaining ROC–AUC as a secondary discrimination metric. Under this mutation-level temporal evaluation, the Essential RF model achieved the strongest numerical performance, with resistant-class F1 of 90.9% and AUC 1.000, although this result is based on only 7 evaluation mutations (6 resistant, 1 susceptible). The Combined RF model achieved resistant-class F1 of 86.8%, resistant-class recall of 80.7%, and AUC 0.735. The Nonessential Logistic model achieved resistant-class F1 of 78.57%, resistant-class recall of 64.71%, and AUC 0.902. Temporal evaluation performance is summarized in Table 2 and Fig. 3B.

**Table 2:** Temporal evaluation on previously uncertain-significance variants in the 2021 WHO catalogue, evaluated against updated WHO 2023 labels. Resistant-class F1 is emphasized because the temporal evaluation set is imbalanced; ninety-five percent Wilson confidence intervals are reported for recall and specificity.

| Dataset | F1<sub>Res</sub> | Precision<sub>Res</sub> | Recall<sub>Res</sub> (95% CI) | Spec. (95% CI) | AUC | Support (R/S) | Incorrect |
| --- | --- | --- | --- | --- | --- | --- | --- |
| Combined | 86.8 | 93.88 | 80.70 (68.66–88.87) | 40.00 (11.76–76.93) | 0.735 | 57/5 | 14 |
| Essential | 90.9 | 100.00 | 83.33 (43.65–96.99) | 100.00 (20.65–100.00) | 1.000 | 6/1 | 1 |
| Nonessential | 78.57 | 100.00 | 64.71 (50.99–76.37) | 100.00 (51.01–100.00) | 0.902 | 51/4 | 18 |

After this temporal evaluation step, we used the Combined RF model as the single deployment model for the full 2023 uncertain-significance variant set. The full 2023 forecasting results, together with the comparison across all three selected models on that deployment pool, are described in Section 2.7.

### 2.5 Comparison to alternative statistical models

Comparison with two alternative statistical models provides an additional robustness check for the multimodal forecasting framework. We compared against a generalized additive model (GAM), which represents nonlinear feature effects with smooth additive functions, and GAQQ, a generative statistical classifier for mixed quantitative and qualitative predictors. On the temporal evaluation benchmark, these alternatives operated differently from the selected multimodal models, with performance that varied substantially across gene strata and often unstable tradeoffs between resistant-class recovery and specificity on the small number of susceptible reclassifications (Supplementary Table S10). In contrast, the selected multimodal Essential RF, Nonessential Logistic, and Combined RF models retained strong resistant-class performance while preserving better balance between resistant and susceptible calls across gene strata.

### 2.6 Interpretability across gene classes

We used interpretability analyses to ask which signals distinguish the Essential, Nonessential, and Combined models, and what the remaining errors reveal about where forecasting is still difficult.

#### 2.6.1 Distinct predictive regimes in the Essential and Nonessential genes

We first examined the five highest-ranked individual features from the best-performing model in each setting. Random forest impurity importances were used for the Essential and Combined models, whereas the Nonessential logistic model was summarized by absolute back-transformed coefficient magnitudes.

We find that the Essential and Combined models are driven by structural and biophysical energy features, with structural context and Rosetta terms consistently near the top. By contrast, the Nonessential model is dominated by protein language model features (Fig. 4A). This pattern is consistent with structural localization playing a stronger role in discrimination for the Essential and Combined settings[28], and our ablation results, whereas the Nonessential setting appears to depend more strongly on broader sequence-derived signal.

**Figure 4:**
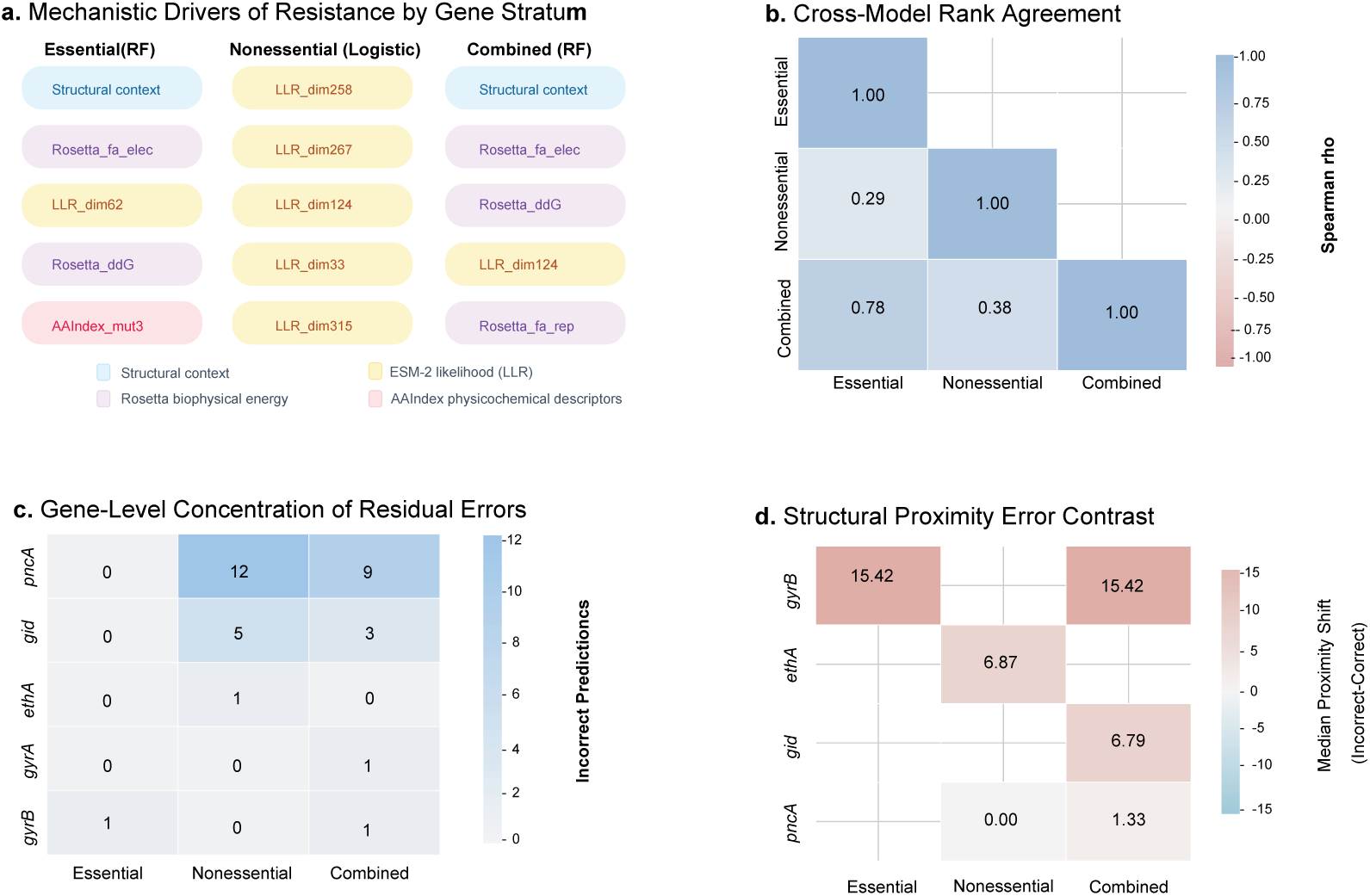
Interpretability across gene classes. (a) Top-ranked features in the selected best models, color-coded by biological feature family and shown by rank rather than raw magnitude to avoid cross-model scale comparisons. (b) Lower-triangle Spearman rank-correlation heatmap comparing feature-importance rank profiles across models. (c) Heatmap of incorrect prediction counts on the 2023 reclassified evaluation set, showing concentration of residual errors in a small number of genes. (d) Median structural-context shift between incorrect and correct calls, where positive values indicate incorrect predictions occur farther from known resistance-associated positions.

We next compared feature-importance rankings across models using Spearman correlation (Fig. 4B). The strongest agreement is between the Essential and Combined models (*ρ* = 0.63, *p* = 6.82 *×*10*^−^*^4^), whereas Essential vs. Nonessential shows no significant alignment (*ρ* = 0.21, *p* = 0.305), and Nonessential vs. Combined is borderline (*ρ* = 0.38, *p* = 0.0587). Thus, pooling the Essential and Nonessential genes does not erase the structural/energetic regime learned in the Essential model; the Combined model remains much closer to Essential than to Nonessential.

#### 2.6.2 Analysis of forecasting errors

We examined whether errors in forecasting were concentrated in specific genes rather than distributed uniformly across the evaluation set (Fig. 4C). By absolute count, most residual errors occur in *pncA* because it contributes the largest number of reclassified variants, but the strongest error enrichment occurs in *gid*. For the Combined RF model, gene identity was significantly associated with error status (*χ*^2^(5) = 12.51, Monte Carlo *p* = 0.031; Section 4.6). Because several gene-level contingency cells had small expected counts, significance was assessed using a fixed-margin Monte Carlo procedure. For the Nonessential Logistic model, the same pattern was even stronger (*χ*^2^(2) = 11.31, Monte Carlo *p* = 0.0036).

Because structural context is the strongest feature family in the Essential and Combined models, we next examined its relationship to incorrect predictions. We separated incorrect predictions into false negatives (FN) and false positives (FP) and compared them with the corresponding correctly classified variants: FN vs. TP for resistant calls and FP vs. TN for susceptible calls. Figure 4D summarizes the resulting median shifts in structural context to known resistance-associated positions. False negatives tend to occur farther from known resistance-associated positions than true positives, and in the Combined model the few false positives occur closer to these anchors than true negatives. This indicates that resistant variants are most often missed when they lie farther from known resistant mutations, whereas susceptible variants called resistant tend to occupy positions unusually close to those anchors.

Overall, the interpretability results support three conclusions. First, the Essential and Combined models rely primarily on structural context and biophysical energy, whereas the Nonessential model follows a distinct protein language model features-dominated regime. Second, the remaining forecasting errors are not distributed uniformly across genes but are concentrated in a small number of loci, with particularly strong enrichment in *gid*. Third, among these residual errors, distance from known resistant positions remains the clearest explanatory signal: variants farther from known anchors are more likely to be missed, whereas the few susceptible variants called resistant tend to lie unusually close to those anchors. Supplementary Fig. S4 provides a deployment-model SHAP view consistent with the same biological interpretation.

### 2.7 Forecasts of future resistance-associated variants

We next applied FARM to the 2023 uncertain-significance mutation set to identify which mutations may in fact be resistance-associated. For deployment-style forecasting, we applied the frozen Combined random forest to all feature-complete 2023 Category 3 uncertain-significance mutations (*n* = 4,525). The model was trained only on labeled 2021 known-effect mutations and used the fixed decision threshold of 0.845 selected from 2021 holdout performance. To contextualize this deployment choice, we also applied all three selected models to the same set of 4,525 feature-complete 2023 Category 3 uncertain-significance mutations. The Essential model classified 886 mutations as resistant (19.6%), the Nonessential model classified 664 (14.7%), and the Combined model classified 696 (15.4%), placing the Combined forecast burden between the higher predicted resistant burden of the Essential model and the lower predicted resistant burden of the Nonessential model (Supplementary Table S8).

Table 3 summarizes, for each gene, the 2021 labeled support, the size of the 2023 uncertain-significance mutation pool, and the number of mutations predicted to be resistance-associated. This comparison helps separate three related quantities: how much direct labeled support a gene had in 2021, how many uncertain mutations are observed for that gene in 2023, and what fraction are predicted to be resistance variants within that pool. Forecasts for genes with limited or absent labeled support should be interpreted with additional caution.

**Table 3:** 2021 training support and forecasted resistant burden per gene for 2023 Category 3 uncertain-significance mutations using the frozen Combined RF model. % Pred. Res. denotes the fraction of mutations classified as resistant at the fixed threshold of 0.845. Support status indicates whether the gene had labeled 2021 training support (labeled), no labeled 2021 support (absent), or only susceptible support in 2021 (susceptible-only). Mean Score is the average random-forest resistance score across all uncertain-significance mutations in that gene. *n_≥0.845_* reports the number of mutations with resistance score at least the deployment threshold.

| Gene | 2021 $n_{res}$ | 2021 $n_{sus}$ | Support Status | 2023 $n_{mut}$ | % Pred. Res. | Mean Score | $n_{\geq 0.845}$ |
| --- | --- | --- | --- | --- | --- | --- | --- |
| *pncA* | 155 | 12 | labeled | 147 | 74.83% | 0.870 | 110 |
| *gid* | 5 | 4 | labeled | 462 | 40.48% | 0.748 | 187 |
| *ethA* | 10 | 0 | labeled | 556 | 32.55% | 0.731 | 181 |
| *rv0678* | 0 | 0 | absent | 154 | 24.03% | 0.720 | 37 |
| *tlyA* | 3 | 0 | labeled | 93 | 20.43% | 0.646 | 19 |
| *atpE* | 0 | 0 | absent | 9 | 11.11% | 0.585 | 1 |
| *rpsL* | 4 | 0 | labeled | 41 | 9.76% | 0.534 | 4 |
| *embB* | 14 | 7 | labeled | 729 | 7.68% | 0.536 | 56 |
| *gyrB* | 5 | 4 | labeled | 290 | 5.86% | 0.519 | 17 |
| *rpoB* | 90 | 7 | labeled | 674 | 5.64% | 0.531 | 38 |
| *katG* | 3 | 2 | labeled | 818 | 4.52% | 0.560 | 37 |
| *ddn* | 0 | 0 | absent | 31 | 3.23% | 0.541 | 1 |
| *gyrA* | 9 | 10 | labeled | 405 | 1.73% | 0.471 | 7 |
| *inhA* | 0 | 1 | susceptible-only | 86 | 1.16% | 0.546 | 1 |
| *rplC* | 0 | 0 | absent | 30 | 0.00% | 0.423 | 0 |

Forecasted resistant burden varies widely across genes. The largest burdens are observed for *pncA*, *gid*, and *ethA*, whereas canonical first-line targets such as *gyrA*, *inhA*, *katG*, *rpoB*, and *embB* show much lower resistant fractions (Table 3). The table also shows that forecast burden is not explained by training support alone: *rv0678* has no direct 2021 labeled support yet 37 of 154 mutations exceed the deployment threshold, whereas *rpoB* has extensive 2021 resistant support but only 38 of 674 uncertain-significance mutations exceed that threshold.

Our model makes predictions even for genes with no mutations in the 2021 labeled set, including *rv0678* and *atpE*. Forecasts in these genes therefore test how the Combined multimodal model, trained across the full labeled 2021 cohort, extrapolates into previously unsupported genes.

Figure 5 maps forecasted residues onto four representative protein structures. The top row shows extrapolation into genes absent from the 2021 labeled training set, whereas the bottom row shows anchor expansion from proteins with known WHO 2021 resistance-associated positions. Orange residues mark known WHO 2021 resistance-associated positions where present, magenta residues mark 2023 uncertain-significance mutations predicted resistant by the frozen Combined RF model, and blue residues mark selected susceptible-side forecasts.

**Figure 5:**
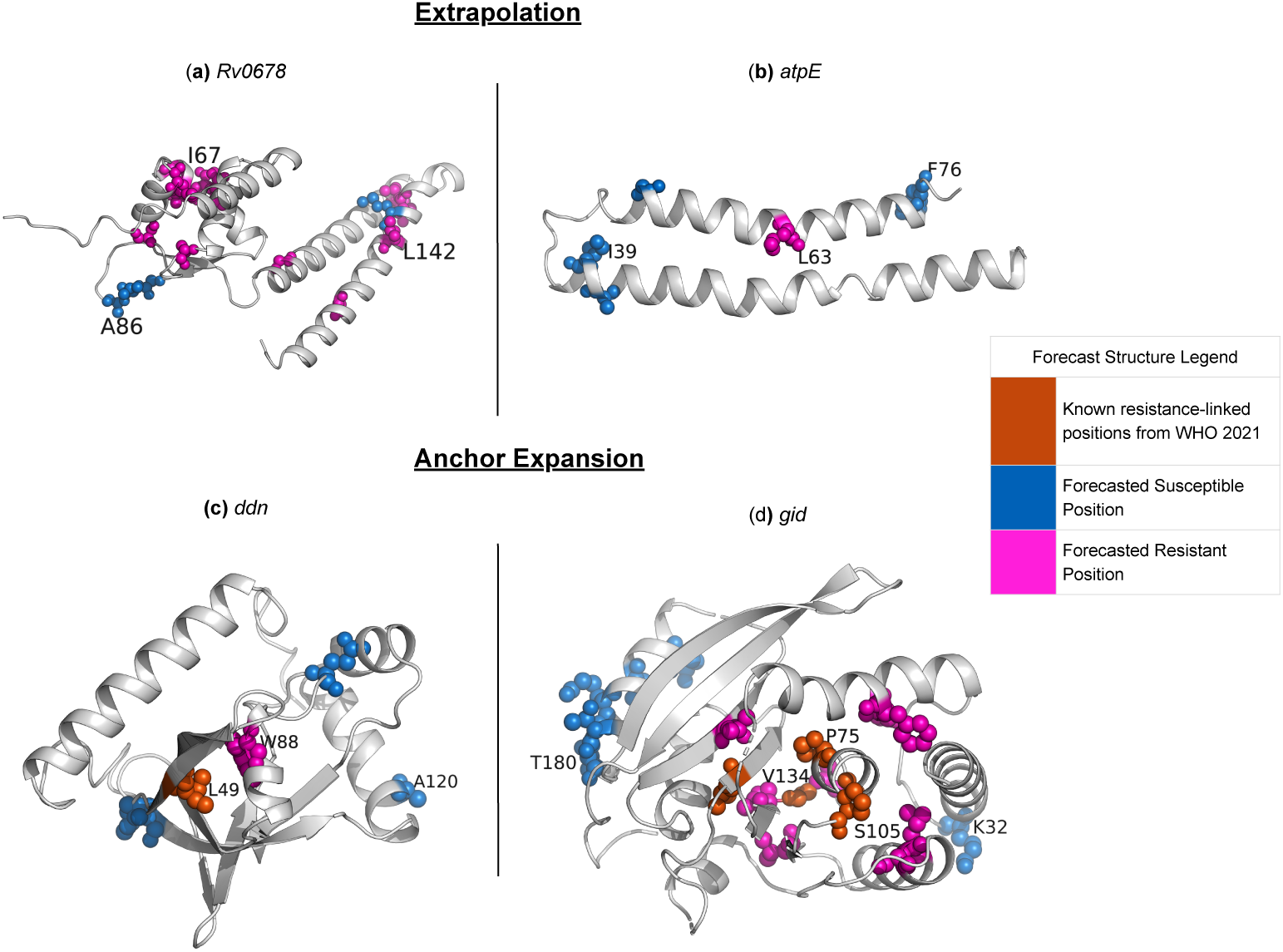
Structural views of forecast behavior in selected proteins encoded by *M. tuberculosis* resistance-associated genes. The top row shows extrapolation into genes absent from the 2021 labeled training set (*rv0678*, *atpE*); the bottom row shows anchor expansion from known WHO 2021 resistance-associated positions (*ddn*, *gid*). Orange residues mark known WHO 2021 resistant positions where present, magenta residues mark 2023 uncertain-significance mutations predicted resistant by the frozen Combined RF model, and blue residues mark selected susceptible-side forecasts. The *rv0678* and *gid* panels show representative hotspot-focused subsets for visualization purposes; full forecast counts are reported in Table 3.

In the top row, the Rv0678 and AtpE proteins show predicted resistant residues concentrated in limited structural regions rather than distributed uniformly across the structure, despite the lack of direct 2021 resistant supervision in the corresponding genes. In the bottom row, newly predicted resistance mutations often occur near known WHO resistance-associated positions. In the Ddn protein, only a single mutation is forecasted to cause antibiotic resistance, with one forecasted resistant site extending from a single known 2021 anchor (Table 3). In the RsmG protein (encoded by the *gid* gene), forecasted resistant positions cluster around known 2021 resistance-associated positions rather than appearing uniformly across the structure. For the *rv0678* gene, 7 forecasted resistant positions overlap functional regions highlighted in prior structural work [35], including the alpha-helix recognition site (62–68) and dimer-stability regions (43–46, 103–108, 135–139, and 151).

*rpoB* provides the clearest negative control: despite strong resistant representation in training, only 5.64% of 2023 Category 3 uncertain-significance *rpoB* mutations are classified as resistant, and the mean resistance score remains close to 0.5. Thus, the model does not simply assign high resistant burden to every gene with substantial historical evidence. Overall, the 2023 forecasts suggest that the Combined model captures a mixture of interpolation and extrapolation behavior. We interpret these predictions as prioritization signals for future catalogue updates and experimental follow-up.

## 3 Discussion

This study introduces FARM, variant-level resistance forecasting using temporally separated WHO catalogue releases. By training on the 2021 WHO catalogue and evaluating against 2023 reclassifications and the 2023 uncertain-significance set, we asked whether multimodal mutation features can prioritize future catalogue updates across genes with potentially different biological constraints. The main result is that they can, but performance is not uniform: the Essential, Nonessential, and Combined models exhibit distinct generalization behavior, and forecasting is most reliable when structurally informative signal or sufficient labeled support is available.

The forecasting results show three complementary behaviors. In genes with stronger 2021 support, the model behaves as a selective interpolator, as illustrated by *rpoB*, where the 2023 uncertain set remains largely on the susceptible side of the decision boundary. In genes such as *pncA*, *gid*, and *ethA*, the model assigns substantially higher resistant burden, consistent with broader nonessential or loss-of-function-like regimes. In genes absent from the 2021 labeled training set, including *Rv0678* and *atpE*, the model still produces structured forecasts when multimodal feature profiles resemble resistant patterns learned elsewhere. The deployment comparison across all three selected models further showed that the Combined model yields an intermediate forecast burden on the full 2023 uncertain set, supporting its use as a single integrated deployment model.

The interpretability analyses provide a biologically coherent explanation for these differences. Essential and Combined predictions are dominated by structural context and energetic perturbation, consistent with constrained mutational neighborhoods in which resistance-conferring substitutions must preserve core protein function. The Nonessential model is dominated by protein language model signal, suggesting that broader sequence-level incompatibility is more informative in genes where many distinct substitutions can disrupt function while remaining compatible with viability. Residual errors are also structured: *gid* remains disproportionately difficult, and false negatives tend to occur farther from known resistance-associated anchors.

The FARM framework has limitations. The labeled training set is small, with substantial class imbalance, and the number of reclassified variants is limited. The reclassification benchmark is also not a random sample of all uncertain variants; it reflects the subset for which enough evidence has accumulated to change WHO status. An additional limitation is the scale of the labeled training set relative to the complexity of modern end-to-end deep learning models. Because the 2021 known-effect cohort contains only a few hundred labeled mutations, we did not train high-capacity architectures such as transformers directly on this task. Instead, we used pretrained protein language model representations as input features within simpler supervised models that are better matched to the available sample size. This design is intended to improve robustness and interpretability under data scarcity, but it also means that the current framework does not fully test whether larger labeled mutation datasets could support more flexible end-to-end models in the future. In addition, the protein language model features are useful predictors but not directly interpretable, and the current random-forest outputs are better interpreted as ranking scores than as calibrated probabilities.

These limitations point to several next steps. Continued temporal validation on future WHO releases should improve calibration and threshold selection, as well as analyses of other mutation catalogues. Leave-one-gene-out or regime-transfer experiments would provide stricter tests of extrapolation into unsupported genes. Broader transfer learning, including pretraining on larger mutation-effect datasets from other organisms, may also improve generalization while preserving the gene-class distinctions observed here. Overall, resistance forecasting from multimodal mutation features appears feasible and informative, but strongly gene-class dependent. We view the forecasts generated by our model as predictions for experimental follow-up and future catalogue curation.

## 4 Methods

### 4.1 Dataset construction and cohort definitions

We constructed the dataset from the 2021 WHO catalogue of *Mycobacterium tuberculosis* resistance-associated variants and generated multimodal sequence/structure features for non-synonymous substitutions.

The WHO catalogue assigns each variant a confidence category describing its association with drug resistance. For supervised learning, we mapped WHO categories 1 and 2 to resistant and categories 4 and 5 to susceptible:

- resistant = {1) Assoc w R, 2) Assoc w R – Interim},
- susceptible = {4) Not assoc w R – Interim, 5) Not assoc w R}.

Starting from the WHO 2021 catalogue, we mapped DNA mutations to non-synonymous amino-acid substitutions and collapsed duplicate entries to one row per gene–mutation, yielding 4,080 unique mutation-level variants. We then retained only feature-complete mutations after quality control, yielding 3,757 model-ready mutations.

These feature-complete mutations were partitioned into distinct downstream analysis sets. First, known-effect variants from WHO categories 1, 2, 4, and 5 were used for binary supervised modeling, yielding 345 labeled mutations (298 resistant, 47 susceptible). Second, Category 3 uncertain-significance variants yielded 3,412 mutation-level uncertain entries that were excluded from supervised training. Among these uncertain 2021 mutations, 62 were reassigned to known-effect categories in the feature-complete 2023 WHO catalogue and therefore provided the temporally separated evaluation set. After model selection and temporal evaluation, the frozen Combined model was also applied to 4,525 mutation-level Category 3 uncertain-significance variants in the 2023 WHO catalogue for deployment-style forecasting.

#### Essentiality stratification

To test whether Essential and Nonessential genes follow different resistance regimes, we stratified the labeled 2021 supervised dataset by operational gene essentiality. We treat mutations in *pncA*, *gid*, and *ethA* as nonessential, and the remaining modeled genes as essential (Table 4). This grouping reflects the observation that *pncA*, *gid*, and *ethA* primarily function as prodrug-activating or accessory loci, where disruption can remain compatible with viability and confer resistance [36, 37], whereas resistance in core targets such as *rpoB* more often requires substitutions that preserve protein function [38–40].

**Table 4:** Labeled dataset grouped by operational gene essentiality.

| Dataset Type | Genes | Total Mutations | Resistant | Susceptible |
| --- | --- | --- | --- | --- |
| Essential | *embB*, *gyrA*, *gyrB*, *inhA*, *katG*,<br>*rpoB*, *rpsL*, *tlyA* | 159 | 128 | 31 |
| Nonessential | *pncA*, *gid*, *ethA* | 186 | 170 | 16 |
| Combined | All above genes | 345 | 298 | 47 |

Each mutation is represented by 25 features spanning biophysical energy, structural context, AAIndex physicochemical descriptors, and protein language model features, as summarized in Table 1. Example feature rows and detailed filtering summaries are provided in the Supplementary Information.

#### 2021–2023 reclassification

For temporal evaluation, we compared the 2021 Category 3 uncertain-significance mutation pool against updated 2023 WHO labels at the mutation level. Among feature-complete matched mutations, 62 uncertain 2021 mutations were reassigned to known-effect categories in 2023 (Table 5).

**Table 5:**
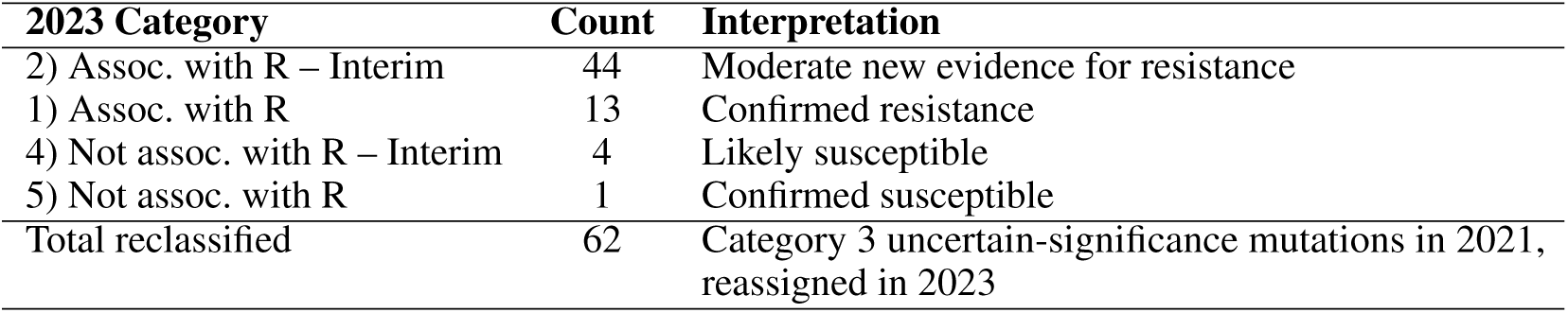
WHO 2021 *→* WHO 2023 outcomes for reclassified uncertain-significance mutations.

| 2023 Category | Count | Interpretation |
| --- | --- | --- |
| 2) Assoc. with R – Interim | 44 | Moderate new evidence for resistance |
| 1) Assoc. with R | 13 | Confirmed resistance |
| 4) Not assoc. with R – Interim | 4 | Likely susceptible |
| 5) Not assoc. with R | 1 | Confirmed susceptible |
| Total reclassified | 62 | Category 3 uncertain-significance mutations in 2021, reassigned in 2023 |

Most reclassified variants were concentrated in a small number of genes. *pncA* accounted for 46 of 62 changes, followed by *gid* (5), *ethA* (4), *gyrB* (4), *katG* (2), and *gyrA* (1), indicating that most new evidence accumulated in nonessential or loss-of-function–prone loci. After model selection and temporal evaluation, we also applied the frozen forecasting model to all 4,525 feature-complete mutation-level Category 3 uncertain-significance mutations in the 2023 WHO catalogue. This all-uncertain 2023 forecast set is distinct from the 62 reclassified variants used for temporal evaluation.

### 4.2 Multimodal feature representation

Each non-synonymous variant was represented using a 25-dimensional multimodal feature vector designed to capture complementary aspects of mutation effect in *Mycobacterium tuberculosis*. The feature set combined four sources of information: (i) physicochemical amino-acid descriptors from AAIndex, (ii) Rosetta-based biophysical-energy and stability terms, (iii) structural context relative to known resistance-associated sites, and (iv) evolutionary plausibility from the ESM-2 protein language model.

AAIndex features were derived from 566 biochemical and physicochemical amino-acid descriptors [41]. Because these descriptors are highly redundant, we applied principal component analysis and retained the first 8 components, which explained 99.5% of the variance [42]. These 8 coordinates represent the physicochemical properties of the mutant residue.

Rosetta-based biophysical-energy features were computed from experimentally derived structures when available and from AlphaFold2 models otherwise [43]. Where needed, missing residues in experimental structures were reconstructed with Modeller [44], and structures were relaxed with Rosetta FastRelax [45]. Mutation effects were then estimated with a PyRosetta workflow similar to our previous study [46], using a 12 Å repacking radius and the REF2015 energy function [47–49]. From this pipeline we retained six biophysical-energy descriptors: `fa_atr, fa_rep, fa_sol, fa_elec, fa_dun`, and ΔΔ*G*. To prevent extreme steric-clash values from dominating the model, energies exceeding 50 REU were squashed with a hyperbolic tangent transformation.

#### Structural Context calculations

Structural context was encoded as the distance from the mutated position to known resistance-associated positions within the same gene. Resistance-associated sites were defined using WHO categories “1) Assoc w R” and “2) Assoc w R – Interim.” We computed both one-dimensional sequence distance and three-dimensional structural context value using EV-couplings distance maps [50], and used the zeroed three-dimensional feature (Prox_3D_zeroed) in the supervised models. The zeroing rule was applied to avoid contradictory per-position labels when multiple catalogue variants mapped to the same residue: when a residue had one or more resistant calls, alternative variants at that same residue were assigned proximity 0 in the zeroed feature. Variants lacking a computable Prox_3D_zeroed value were excluded from downstream forecasting.

Evolutionary plausibility was summarized using ESM-2 (150M parameters) in a masked-marginal scoring framework [51, 52]. For each missense mutation, we computed the scalar log-likelihood ratio comparing the mutant and wild-type residues in sequence context, and additionally derived a mutation-specific latent contribution vector from the model representation at the mutated position. We retained the 10 latent coordinates with the largest mean absolute magnitude across mutations, forming the protein language model feature set. The final supervised feature matrix therefore comprised 8 AAIndex mutant descriptors, 6 Rosetta-based energetic terms, 1 structural context feature, and 10 protein language model features.

### 4.3 Training design and temporal forecasting setup

Our goal was to train on known-effect mutations and then forecast which uncertain-significance mutations may later be recognized as resistance-conferring. Supervised learning used mutation-level entries derived from the 2021 WHO catalogue: known-effect variants from categories 1, 2, 4, and 5 were collapsed to one row per labeled gene–mutation entry, yielding 345 labeled mutations (298 resistant and 47 susceptible), while uncertain-significance mutations were excluded from fitting. These labeled mutations were divided into an 80% training split and a 20% holdout split for model selection and evaluation.

We considered three gene-grouping settings motivated by biological differences between gene strata: Essential, Nonessential, and Combined. The Essential and Combined settings were best captured by random forests, whereas the Nonessential setting favored logistic regression. We therefore treated model comparison as a selection step rather than a claim that one model family was universally optimal. The final deployment-style forecast used the frozen Combined random forest, because it provided the strongest overall balance between discrimination and sensitivity in the temporally realistic evaluation.

After training on the 2021 known-effect mutations, we evaluated forecasting in two stages. First, we applied the models to the 2021 uncertain-significance mutations pool, collapsed to one row per gene–mutation, and compared predictions against the subset of uncertain-significance mutations that could be matched to the feature-complete 2023 WHO catalogue. Under this mutation-level temporal evaluation benchmark, 62 uncertain 2021 mutations were reassigned to known-effect categories in 2023. This temporal evaluation therefore targets future catalogue reclassification rather than a random sample of all future biological ground truth. Second, after model selection, we applied the frozen Combined model to all 4,525 feature-complete uncertain-significance mutations in the 2023 WHO catalogue. This design mimics a real-world forecasting scenario in which predictions must be made before future catalogue revisions are known.

### 4.4 Prediction models and threshold selection

Resistance prediction was framed as a binary classification task (resistant vs. susceptible) using the 25-dimensional multimodal representation described above. For the Nonessential setting, we used an sklearn pipeline consisting of `StandardScaler` followed by `LogisticRegression` with `solver=liblinear`, `max_iter=1000`, and `class_weight=balanced` [53]. For the Essential and Combined settings, we used RandomForestClassifier with n_estimators=200, max_depth=None, class_weight=balanced_subsample, and random_state=42. All remaining parameters were left at library defaults.

To stabilize classification thresholds under class imbalance, thresholds were selected using repeated stratified cross-validation on the training split only (`RepeatedStratifiedKFold`, 5 folds, 3 repeats, `random_state=42`). For each fold, we identified the threshold maximizing Youden’s index, and the final operating threshold was set to the median across folds and repeats. These fixed thresholds were then applied once to the holdout data and subsequently reused without retuning for the temporal forecasting analyses.

We report ROC–AUC, sensitivity, specificity, and related thresholded metrics on both the 2021 holdout set and the temporally separated 2023 temporal evaluation set. For the random-forest models, global feature importance was quantified by mean decrease in Gini impurity; for the Nonessential logistic model, interpretability used the coefficient-based summaries (Section 2.6).

### 4.5 Comparison with alternative statistical models

To test whether the forecasting results depended strongly on the chosen classifier family, we additionally compared the random forest with two alternative statistical frameworks: a generalized additive model (GAM) [54] and the Generative Approach for Quantitative and Qualitative Responses (GAQQ) [55]. These models were used as robustness checks rather than as the primary forecasting pipeline. Their full formulations and performance summaries under the mutation-level benchmark are reported in the Supplementary Results.

### 4.6 Error analysis

Because false negatives and false positives have different biological implications in resistance prediction, we performed targeted residual-error analyses after model fitting. These included gene-level enrichment of misclassifications, group-wise summaries of remaining error signal, and false-negative/false-positive comparisons centered on structural context. These analyses were designed to clarify not only which models performed best, but also where and why the remaining forecasting errors arose.

### 4.7 Statistical analysis

All reported sample sizes (*n*) correspond to the number of mutation-level entries included in the relevant analysis. Model discrimination was summarized using ROC–AUC, and thresholded performance was summarized using accuracy, sensitivity, specificity, precision, recall, and F1 score as appropriate. Ninety-five percent confidence intervals for sensitivity and specificity were computed using Wilson intervals. For gene-level enrichment of forecasting errors, we analyzed contingency tables using chi-square statistics. Because several expected cell counts were small, significance was assessed using a fixed-margin Monte Carlo procedure rather than relying only on the asymptotic chi-square approximation.

## Code availability

The code used for data preparation, model training, temporal evaluation, and deployment-style forecasting is available at https://github.com/SAGE-Lab-UMass/resistance_forecast.

## Data availability

The WHO 2021 and WHO 2023 tuberculosis mutation catalogues are available from the publicly available sources provided by World Health Organization [34, 6]. A curated manuscript-facing data package, including the source catalogue metadata and the training data used for modeling, is available through the same GitHub repository at https://github.com/SAGE-Lab-UMass/resistance_ forecast. The full set of 4,525 Combined-model forecasts for feature-complete WHO 2023 Category 3 uncertain-significance mutations is provided as Supplementary Data 1.

## Author contributions

A.G.G., M.T., and J.C. conceived the study. M.T. and S.B. curated the data. M.T. developed the methodology and software. M.T. and Y.W. performed the formal analysis. All authors interpreted the results. M.T. and S.B. prepared the visualizations. A.G.G., J.C., and L.K. supervised the study. All authors wrote and edited the manuscript.

## Materials & correspondence

Correspondence and requests for materials should be addressed to M.T. or A.G.G.

## Acknowledgements

This work was supported by a University of Massachusetts Interdisciplinary Research Grant to AGG, JC, and LK. We thank the members of the UMass SAGE lab for helpful discussion of the work. This work utilized resources from Unity, a collaborative, multi-institutional high-performance computing cluster managed by UMass Amherst Research Computing and Data.

## SUPPLEMENTARY MATERIALS

### List of supplementary figures

Fig. S1 Distribution of mutations across eleven essential and nonessential proteins listed in the WHO 2021 catalog.

Fig. S2 Distribution of mutations across eleven essential and nonessential proteins listed in the WHO 2023 catalog.

Fig. S3 Model-family comparison on the 2021 holdout set across the Essential, Nonessential, and Combined dataset configurations.

Fig. S4 SHAP beeswarm summary for the Combined random forest deployment model.

### List of supplementary tables

Table S1 WHO confidence categories used in the 2021/2023 forecasting workflow.

Table S2 Example rows illustrating the feature representation for individual mutations.

Table S3 Summary of 2021 catalogue counts at each filtering stage.

Table S4 Cohort breakdown for supervised fitting, temporal evaluation, and deployment-style forecasting.

Table S5 Training/holdout performance per dataset/model.

Table S6 Holdout performance of structural context-only baselines across datasets and model families.

Table S7 Holdout performance of feature-group ablations for the best-performing model per dataset.

Table S8 Full 2023 uncertain-significance mutation-set comparison across the three selected deployment models.

Table S9 Proteins-of-interest summary for the structural forecast panels in the main text.

Table S10 Temporal evaluation performance of alternative statistical models (GAM and GAQQ) on Category 3 uncertain-significance mutations.

Table S11 Feature importances and ranks across Essential, Nonessential, and Combined models.

### A Supplement Overview

This Supplement contains additional dataset-composition and filtering summaries for the 2021 and 2023 WHO catalogues, full temporal-evaluation model-comparison and ablation results, temporally separated Category 3 uncertain-significance evaluation summaries, interpretability analyses for the Combined deployment model and cross-model comparisons, and extended benchmarking against alternative statistical models.

### B Dataset Supplemental

This section provides compact reference tables for the mutation-level cohort definitions used in the manuscript, including the WHO confidence categories, the 2021 filtering summary, and the final supervised, temporal-evaluation, and deployment cohorts.

Table S1 summarizes the WHO confidence categories used throughout the manuscript.

**Table S1:**
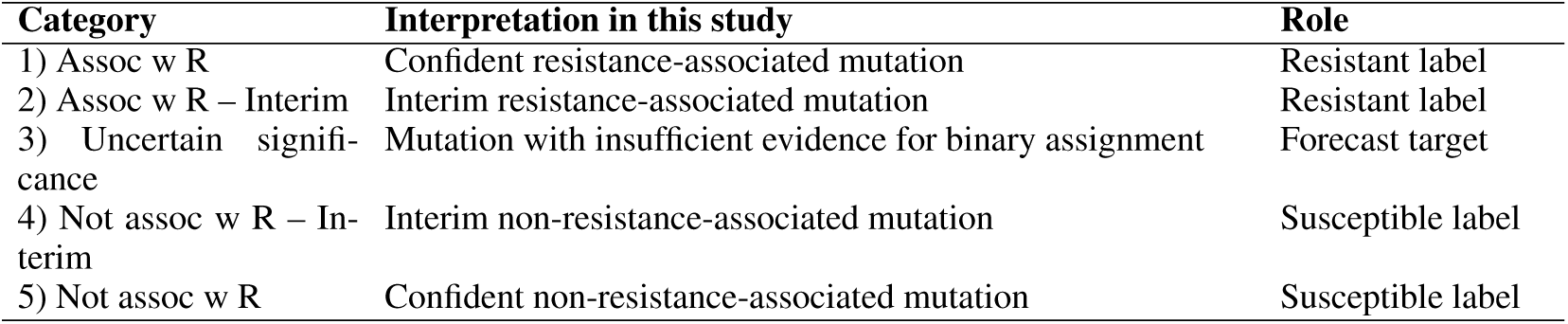
WHO confidence categories used in the 2021/2023 forecasting workflow.

| Category | Interpretation in this study | Role |
| --- | --- | --- |
| 1) Assoc w R | Confident resistance-associated mutation | Resistant label |
| 2) Assoc w R – Interim | Interim resistance-associated mutation | Resistant label |
| 3) Uncertain significance | Mutation with insufficient evidence for binary assignment | Forecast target |
| 4) Not assoc w R – Interim | Interim non-resistance-associated mutation | Susceptible label |
| 5) Not assoc w R | Confident non-resistance-associated mutation | Susceptible label |

Table S2 provides a compact illustration of representative feature-matrix rows.

**Table S2:** Example rows illustrating the feature representation for individual mutations. Only a subset of columns is shown for clarity.

| gene | drug | confidence | phenotype | mutation | fa_atr | AAIndex_mut3 |
| --- | --- | --- | --- | --- | --- | --- |
| embB | EMB | 2) Assoc w R–Interim | Resistant | L74R | -2.640 | 1.268 |
| embB | EMB | 5) Not assoc w R | Susceptible | Q139H | 0.343 | 4.664 |
| embB | EMB | 4) Not assoc w R–Interim | Susceptible | G156C | -5.335 | 6.016 |

Table S3 summarizes the key filtering stages for the 2021 mutation-level cohort.

**Table S3:** Summary of 2021 catalogue counts at each filtering stage under the mutation-level workflow.

| Category | Count | Notes |
| --- | --- | --- |
| Unique mutation-level non-synonymous variants | 4,080 | After mapping DNA mutations to amino-acid substitutions and merging duplicate catalogue entries to one gene–mutation record |
| Feature-complete model-ready mutations | 3,757 | Retained after quality control filtering for complete feature availability |
| Known effect (1/2/4/5) | 345 | Mutation-level supervised learning set |
| Category 3 uncertain-significance | 3,412 | Mutation-level temporal evaluation pool |

Table S4 is showing the four cohorts used in the manuscript: the 2021 known-effect supervised set, the 2021 Category 3 uncertain-significance temporal-evaluation pool, the 2023 reclassified evaluation subset, and the 2023 all-uncertain deployment mutation set.

**Table S4:** Cohort breakdown for supervised fitting, temporal evaluation, and deployment-style forecasting.

| Cohort | Count | Use in this study |
| --- | --- | --- |
| 2021 known-effect labeled mutations (categories 1/2/4/5) | 345 | Supervised fitting and holdout model comparison |
| 2021 Category 3 uncertain-significance mutations | 3,412 | Temporal evaluation pool |
| 2021 Category 3 uncertain-significance mutations reclassified by 2023 | 62 | Temporally separated evaluation subset |
| 2023 feature-complete Category 3 uncertain-significance mutations | 4,525 | Deployment-style forecasting target set for the frozen Combined RF |

### C Result supplemental

Before selecting a final model for each dataset configuration, we compared both logistic-regression and random-forest classifiers across the Essential, Nonessential, and Combined settings. Supplementary Table S5 reports the full holdout and cross-validation results for all model–dataset combinations, showing that model selection was comparative rather than arbitrary.

**Figure S1:**
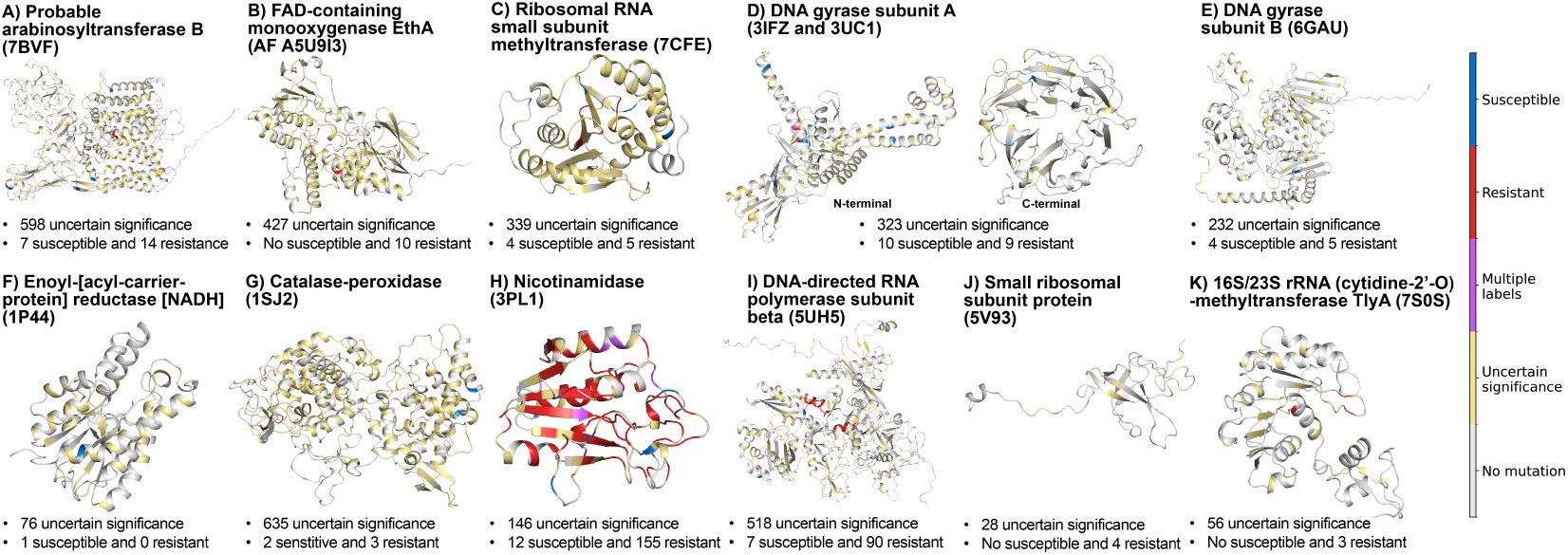
Supplementary: Distribution of mutations across eleven essential and nonessential proteins listed in the WHO 2021 catalog. Colors represent different mutation categories at specific positions (blue: susceptible, red: resistant, purple: both susceptible and resistant, yellow: unknown significance), and gray indicates no mutation detected.

**Figure S2:**
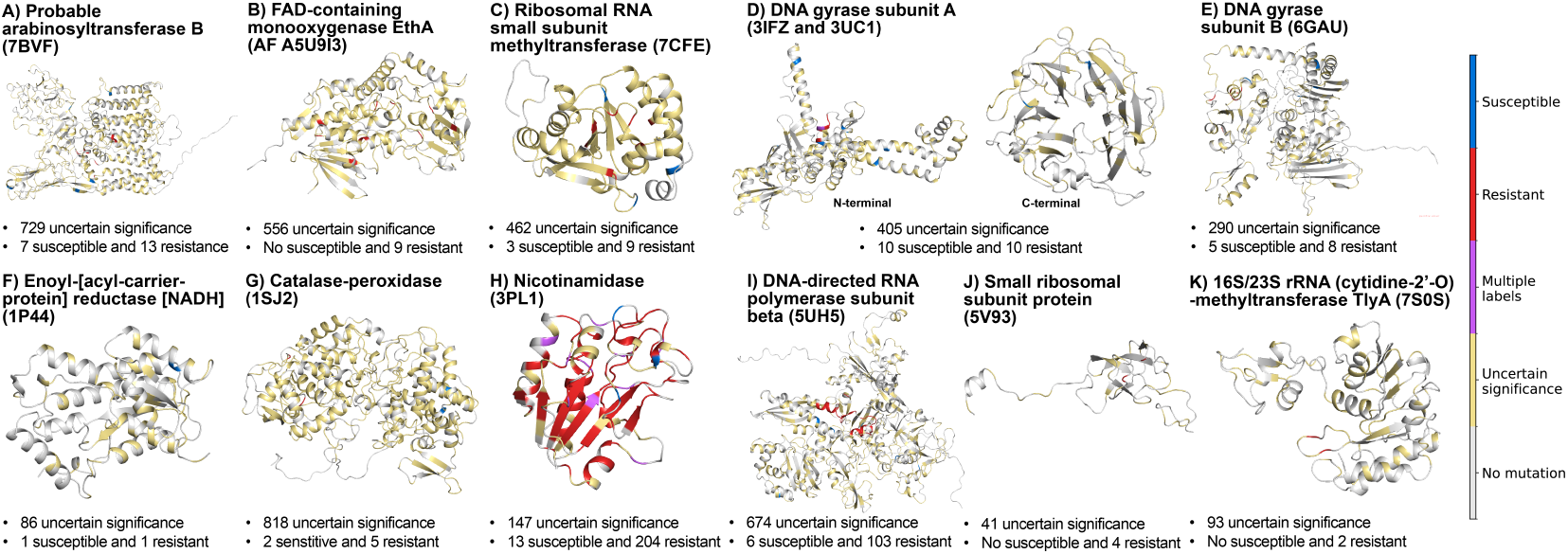
Supplementary: Distribution of mutations across eleven essential and nonessential proteins listed in the WHO 2023 catalog. Colors represent different mutation categories at specific positions (blue: susceptible, red: resistant, purple: both susceptible and resistant, yellow: unknown significance), and gray indicates no mutation detected.

**Figure S3:**
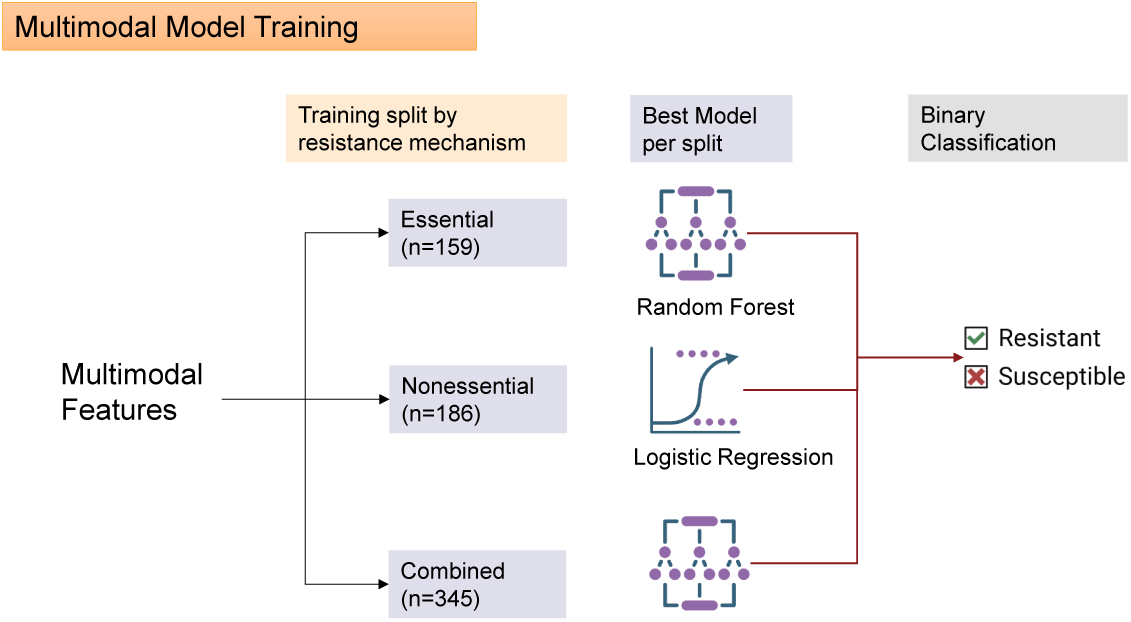
Model-family comparison on the 2021 holdout set across the Essential, Nonessential, and Combined dataset configurations.

**Table S5:** Training/holdout performance per dataset/model under the mutation-level workflow.

| Dataset | Model | Mode | AUC | Accuracy | Sensitivity | Specificity | Threshold |
| --- | --- | --- | --- | --- | --- | --- | --- |
| Essential | Logistic | holdout | 0.808 | 0.875 | 0.923 | 0.667 | 0.706 |
| Essential | Logistic | cv | 0.884 | 0.913 | 0.946 | 0.771 | 0.601 |
| Essential | RF | holdout | 0.843 | 0.906 | 0.923 | 0.833 | 0.810 |
| Essential | RF | cv | 0.938 | 0.950 | 0.954 | 0.938 | 0.730 |
| Nonessential | Logistic | holdout | 0.943 | 0.842 | 0.829 | 1.000 | 0.814 |
| Nonessential | Logistic | cv | 0.840 | 0.882 | 0.912 | 0.583 | 0.374 |
| Nonessential | RF | holdout | 0.671 | 0.789 | 0.800 | 0.667 | 0.935 |
| Nonessential | RF | cv | 0.753 | 0.822 | 0.859 | 0.433 | 0.860 |
| Combined | Logistic | holdout | 0.894 | 0.928 | 0.933 | 0.889 | 0.644 |
| Combined | Logistic | cv | 0.913 | 0.864 | 0.882 | 0.749 | 0.562 |
| Combined | RF | holdout | 0.920 | 0.942 | 0.983 | 0.667 | 0.845 |
| Combined | RF | cv | 0.931 | 0.887 | 0.916 | 0.702 | 0.780 |

**Table S6:** Holdout performance of structural context-only baselines.

| Dataset | Model | AUC | Accuracy | Sensitivity | Specificity | Threshold |
| --- | --- | --- | --- | --- | --- | --- |
| Essential | Logistic | 0.968 | 0.906 | 0.923 | 0.833 | 0.757 |
| Essential | RF | 0.811 | 0.906 | 0.923 | 0.833 | 0.948 |
| Nonessential | Logistic | 0.757 | 0.868 | 0.886 | 0.667 | 0.519 |
| Nonessential | RF | 0.814 | 0.868 | 0.886 | 0.667 | 0.699 |
| Combined | Logistic | 0.863 | 0.899 | 0.933 | 0.667 | 0.640 |
| Combined | RF | 0.788 | 0.928 | 0.967 | 0.667 | 0.750 |

**Table S7:** Holdout performance of feature-group ablations for the best-performing model per dataset under the mutation-level workflow. Feature-group names are reported using the manuscript conventions: protein language model features, biophysical energy, structural context, and AAIndex physicochemical descriptors.

| Dataset | Ablation | AUC | Sens. | Spec. | Thresh. |
| --- | --- | --- | --- | --- | --- |
| Essential | keep all | 0.843 | 0.923 | 0.833 | 0.810 |
| Essential | drop Protein language model features | 0.827 | 0.923 | 0.833 | 0.775 |
| Essential | drop Biophysical Energy | 0.833 | 0.962 | 0.833 | 0.830 |
| Essential | drop Structural Context | 0.760 | 0.385 | 0.833 | 0.835 |
| Essential | drop AAIndex physicochemical descriptors | 0.830 | 0.923 | 0.833 | 0.835 |
| Essential | drop Protein language model features + Biophysical Energy | 0.885 | 0.962 | 0.833 | 0.691 |
| Essential | drop Protein language model features + Biophysical Energy + Structural Context | 0.673 | 0.346 | 0.833 | 0.926 |
| Essential | drop Protein language model features + Biophysical Energy + AAIndex physicochemical descriptors | 0.811 | 0.923 | 0.833 | 0.948 |
| Nonessential | keep all | 0.943 | 0.829 | 1.000 | 0.814 |
| Nonessential | drop Protein language model features | 0.914 | 0.686 | 1.000 | 0.693 |
| Nonessential | drop Biophysical Energy | 0.876 | 0.600 | 1.000 | 0.871 |
| Nonessential | drop Structural Context | 0.914 | 0.743 | 1.000 | 0.873 |
| Nonessential | drop AAIndex physicochemical descriptors | 0.895 | 0.857 | 0.667 | 0.583 |
| Nonessential | drop Protein language model features + Biophysical Energy | 0.833 | 0.457 | 1.000 | 0.700 |
| Nonessential | drop Protein language model features + Biophysical Energy + Structural Context | 0.624 | 0.371 | 1.000 | 0.647 |
| Nonessential | drop Protein language model features + Biophysical Energy + AAIndex physicochemical descriptors | 0.757 | 0.886 | 0.667 | 0.519 |
| Combined | keep all | 0.920 | 0.983 | 0.667 | 0.845 |
| Combined | drop Protein language model features | 0.900 | 0.983 | 0.667 | 0.845 |
| Combined | drop Biophysical Energy | 0.916 | 0.950 | 0.778 | 0.835 |
| Combined | drop Structural Context | 0.746 | 0.767 | 0.556 | 0.855 |
| Combined | drop AAIndex physicochemical descriptors | 0.818 | 0.983 | 0.667 | 0.825 |
| Combined | drop Protein language model features + Biophysical Energy | 0.859 | 0.933 | 0.556 | 0.722 |

Table S7: Supplementary: Holdout performance of feature-group ablations for the best-performing model per dataset under the mutation-level workflow.
| Dataset | Ablation | AUC | Sens. | Spec. | Thresh. |
| --- | --- | --- | --- | --- | --- |
| Combined | drop Protein language model features + Biophysical Energy + Structural Context | 0.511 | 0.783 | 0.222 | 0.507 |
| Combined | drop Protein language model features + Biophysical Energy + AAIndex physicochemical descriptors | 0.788 | 0.967 | 0.667 | 0.750 |

**Table S8:** Full 2023 uncertain-significance mutation-set comparison across the three selected deployment models. Each selected model was applied to the same set of 4,525 feature-complete WHO 2023 uncertain-significance mutations using its training-derived fixed threshold.

| Model setting | *n* mutations | Pred. resistant fraction | Pred. resistant count | Model key | Threshold |
| --- | --- | --- | --- | --- | --- |
| Essential | 4525 | 0.1960 | 886 | Essential_rf | 0.810 |
| Nonessential | 4525 | 0.1470 | 664 | Nonessential_logistic | 0.814 |
| Combined | 4525 | 0.1540 | 696 | Combined_rf | 0.845 |

#### Performance Comparison of Alternative Statistical Models

Table S10 summarizes the temporal evaluation performance of two alternative statistical models, GAM and GAQQ, under the revised mutation-level workflow. These models were evaluated on 2021 Category 3 uncertain-significance mutations reclassified in 2023 using the same mutation-level temporal evaluation benchmark applied in the revised manuscript pipeline. Across datasets, these alternatives show unstable operating behavior: some settings achieve high resistant sensitivity, but often at the cost of weak susceptible specificity. Overall, these results support the stronger balance achieved by the primary multimodal models.

Table S10 summarizes the temporal evaluation performance of two alternative statistical models, GAM and GAQQ, evaluated on Category 3 uncertain-significance mutations using the same mutation-level benchmark as the primary manuscript models. Results are reported separately for the Essential, Nonessential, and Combined gene sets. Across datasets, these alternatives do not show a consistent operating regime. GAM performs well in the small Essential set (AUC 100.0, sensitivity 83.3%, specificity 100.0) but drops to weaker discrimination in the Nonessential and Combined settings. GAQQ retains moderate resistant sensitivity in the Nonessential set (76.5%) but collapses in the Combined setting, where sensitivity falls to 1.8% despite perfect specificity.

These shifts should be interpreted in the context of the temporal evaluation sets, which are strongly resistant-enriched and contain very few susceptible reclassifications. Under this benchmark, the multimodal manuscript models remain preferable because they provide a more stable balance between resistant recovery and susceptible discrimination across gene strata than either GAM or GAQQ alone.

**Table S9:** Proteins-of-interest summary for the structural forecast panels in the main text. Values are taken from the frozen Combined RF forecasts on the 2023 Category 3 uncertain-significance mutation set. Residue positions correspond to the representative positions highlighted in Figure 5; the complete per-gene forecast burden is reported in the main-text forecast table.

| Gene | Panel type | 2021 support | 2023 $n$ | % Pred. Res. | Highlighted resistant / candidate positions | Highlighted susceptible-side positions |
| --- | --- | --- | --- | --- | --- | --- |
| *rv0678* | Extrapolation | – / – | 154 | 24.03% | 67, 142 | 86 |
| *atpE* | Extrapolation | – / – | 9 | 11.11% | 63 | 39, 76 |
| *ddn* | Anchor expansion | – / – | 31 | 3.23% | anchor 49; forecast 88 | 120 |
| *gid* | Anchor expansion | 5 / 4 | 462 | 40.48% | anchors 75, 134; forecast 105 | 32, 180 |

**Table S10:** Temporal evaluation performance of alternative statistical models (GAM and GAQQ) on Category 3 uncertain-significance mutations reclassified between 2021 and 2023.

| Dataset | Model | AUC | Sens (R) | Spec (S) | Support R | Support S | Incorrect |
| --- | --- | --- | --- | --- | --- | --- | --- |
| Essential | GAM | 100.00 | 83.3 | 100.0 | 6 | 1 | 1 |
| Nonessential | GAM | 52.94 | 33.3 | 75.0 | 51 | 4 | 35 |
| Combined | GAM | 55.79 | 86.0 | 40.0 | 57 | 5 | 11 |
| Essential | GAQQ | 100.00 | 33.3 | 100.0 | 6 | 1 | 4 |
| Nonessential | GAQQ | 55.88 | 76.5 | 0.0 | 51 | 4 | 16 |
| Combined | GAQQ | 50.53 | 1.8 | 100.0 | 57 | 5 | 56 |

### D Interpretability

#### Feature-importance ranking across the three selected models

**Table S11:**
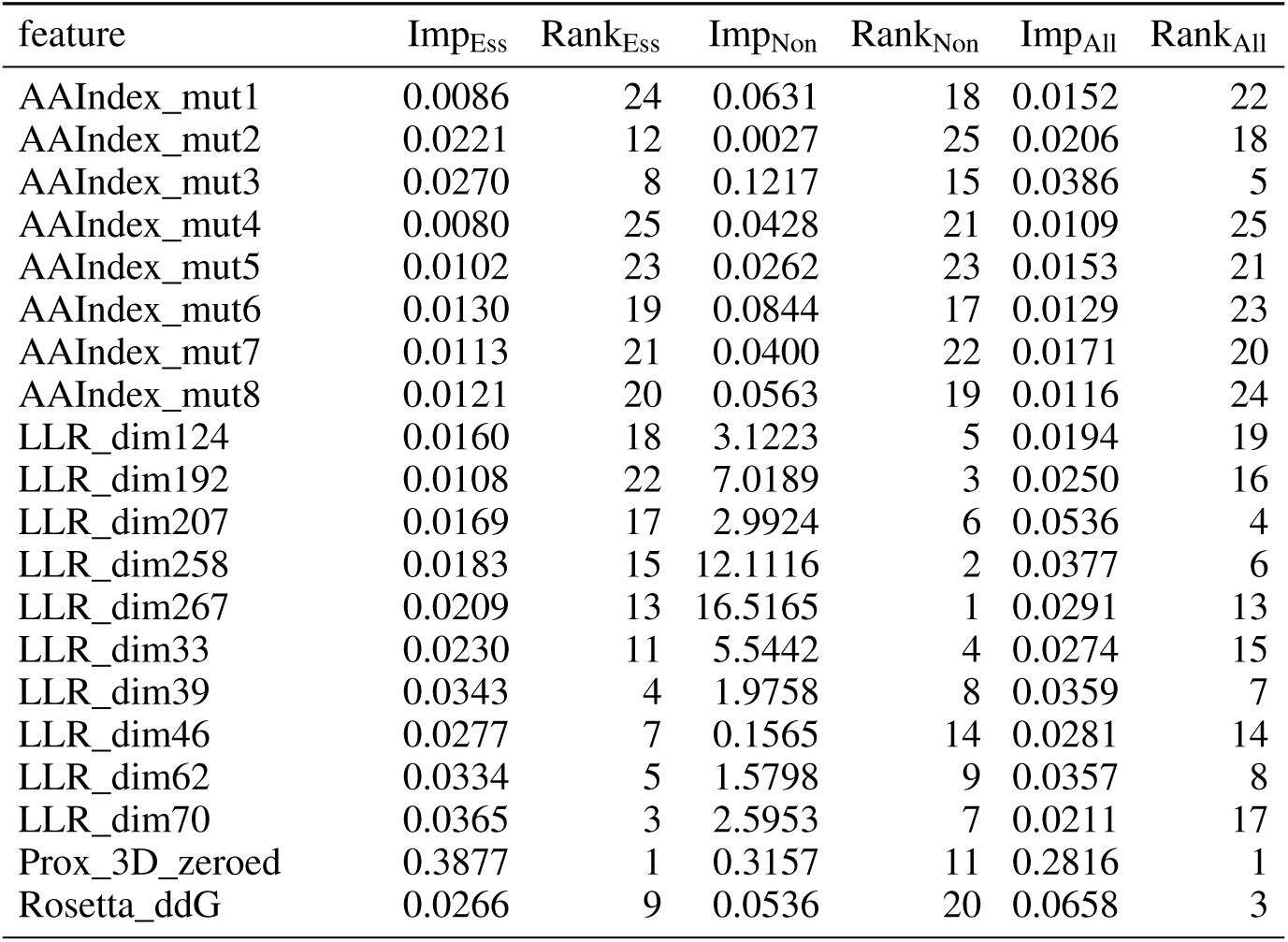
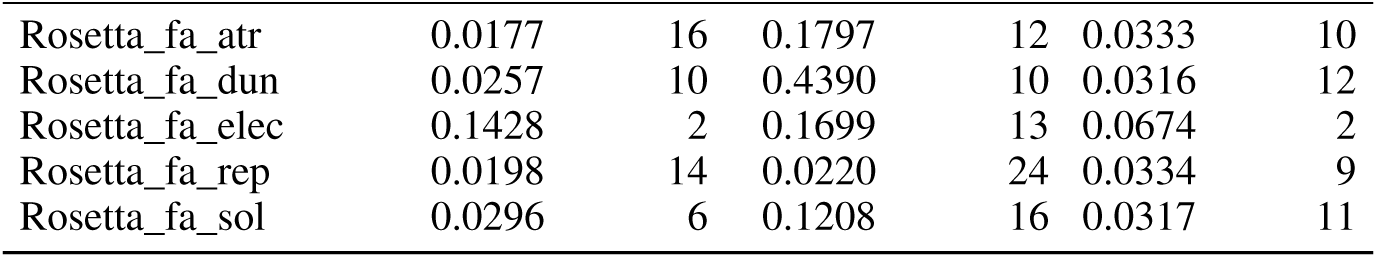
Feature importances and ranks across the three selected models. Importance (Imp) and rank are reported for the Essential, Nonessential, and Combined settings.

| feature | Imp<sub>Ess</sub> | Rank<sub>Ess</sub> | Imp<sub>Non</sub> | Rank<sub>Non</sub> | Imp<sub>All</sub> | Rank<sub>All</sub> |
| --- | --- | --- | --- | --- | --- | --- |
| AAIndex_mut1 | 0.0086 | 24 | 0.0631 | 18 | 0.0152 | 22 |
| AAIndex_mut2 | 0.0221 | 12 | 0.0027 | 25 | 0.0206 | 18 |
| AAIndex_mut3 | 0.0270 | 8 | 0.1217 | 15 | 0.0386 | 5 |
| AAIndex_mut4 | 0.0080 | 25 | 0.0428 | 21 | 0.0109 | 25 |
| AAIndex_mut5 | 0.0102 | 23 | 0.0262 | 23 | 0.0153 | 21 |
| AAIndex_mut6 | 0.0130 | 19 | 0.0844 | 17 | 0.0129 | 23 |
| AAIndex_mut7 | 0.0113 | 21 | 0.0400 | 22 | 0.0171 | 20 |
| AAIndex_mut8 | 0.0121 | 20 | 0.0563 | 19 | 0.0116 | 24 |
| LLR_dim124 | 0.0160 | 18 | 3.1223 | 5 | 0.0194 | 19 |
| LLR_dim192 | 0.0108 | 22 | 7.0189 | 3 | 0.0250 | 16 |
| LLR_dim207 | 0.0169 | 17 | 2.9924 | 6 | 0.0536 | 4 |
| LLR_dim258 | 0.0183 | 15 | 12.1116 | 2 | 0.0377 | 6 |
| LLR_dim267 | 0.0209 | 13 | 16.5165 | 1 | 0.0291 | 13 |
| LLR_dim33 | 0.0230 | 11 | 5.5442 | 4 | 0.0274 | 15 |
| LLR_dim39 | 0.0343 | 4 | 1.9758 | 8 | 0.0359 | 7 |
| LLR_dim46 | 0.0277 | 7 | 0.1565 | 14 | 0.0281 | 14 |
| LLR_dim62 | 0.0334 | 5 | 1.5798 | 9 | 0.0357 | 8 |
| LLR_dim70 | 0.0365 | 3 | 2.5953 | 7 | 0.0211 | 17 |
| Prox_3D_zeroed | 0.3877 | 1 | 0.3157 | 11 | 0.2816 | 1 |
| Rosetta_ddG | 0.0266 | 9 | 0.0536 | 20 | 0.0658 | 3 |

Table S11: Feature importances and ranks across models (continued).
| feature | Imp<sub>Ess</sub> | Rank<sub>Ess</sub> | Imp<sub>Non</sub> | Rank<sub>Non</sub> | Imp<sub>All</sub> | Rank<sub>All</sub> |
| --- | --- | --- | --- | --- | --- | --- |
| Rosetta_fa_atr | 0.0177 | 16 | 0.1797 | 12 | 0.0333 | 10 |
| Rosetta_fa_dun | 0.0257 | 10 | 0.4390 | 10 | 0.0316 | 12 |
| Rosetta_fa_elec | 0.1428 | 2 | 0.1699 | 13 | 0.0674 | 2 |
| Rosetta_fa_rep | 0.0198 | 14 | 0.0220 | 24 | 0.0334 | 9 |
| Rosetta_fa_sol | 0.0296 | 6 | 0.1208 | 16 | 0.0317 | 11 |

**Figure S4:**
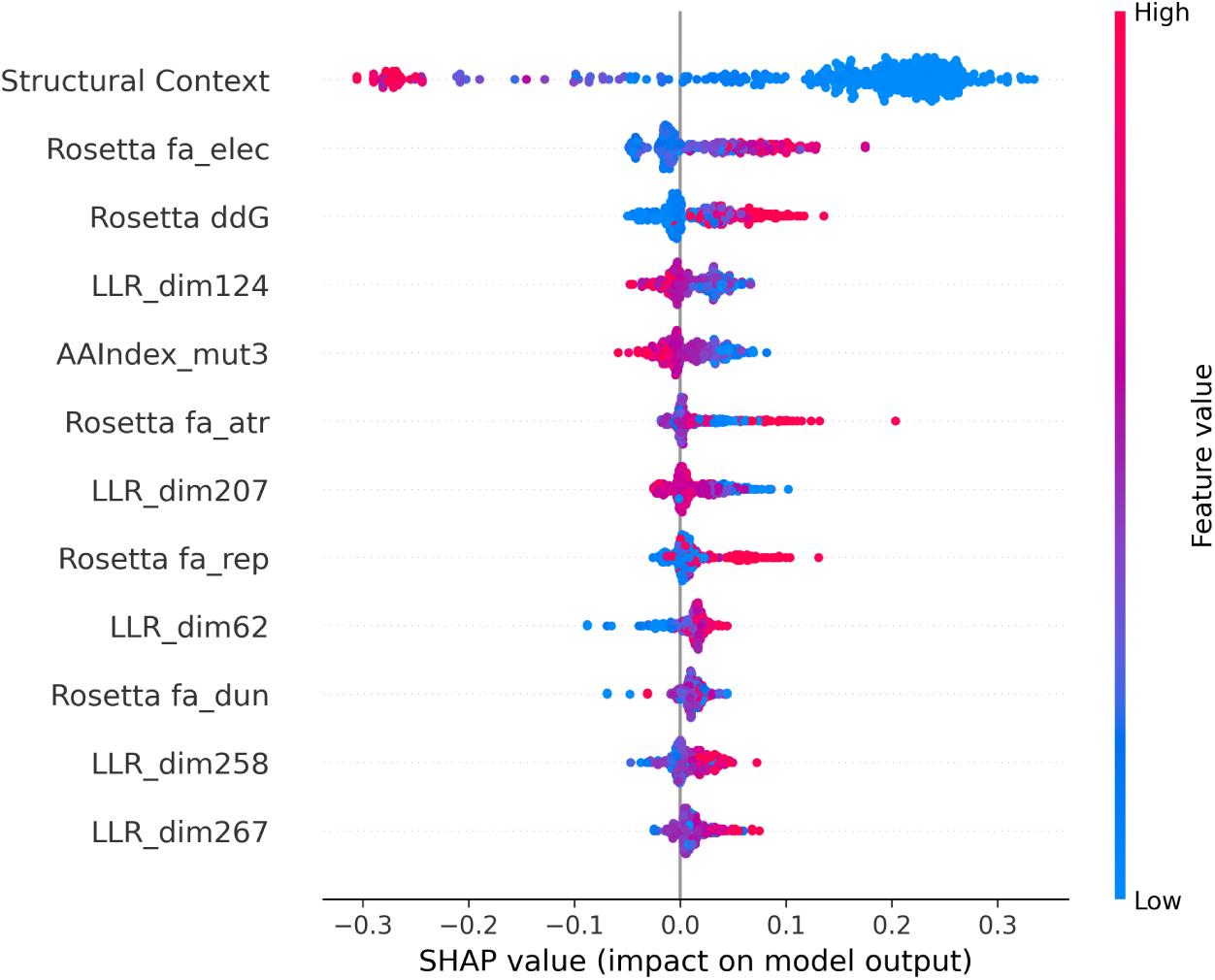
SHAP beeswarm summary for the refit Combined RF model. Each point represents one labeled 2021 mutation from the Combined training set, positioned by its SHAP value for the resistant class. Color indicates the underlying feature value.

